# Rational Expectations and Kinematic Information in Coordination Games

**DOI:** 10.1101/2025.08.01.668238

**Authors:** Martina Fanghella, Camilla F. Colombo, Maria Teresa Pascarelli, Guido Barchiesi, Marco Rabuffetti, Maurizio Ferrarin, Francesco Guala, Corrado Sinigaglia

## Abstract

Successful coordination often requires integrating strategic reasoning with real-time observations of others’ actions, yet how humans resolve conflicts between these information sources remains unclear. This study aimed to fill this gap by examining how people coordinate in a strategic game when observing partial kinematic information from their partner’s actions. Participants played a HI-LO game with a virtual partner, choosing payoffs based on grasping movements toward invisible large and small targets. Hand movements were presented as schematic animations, with partners grasping targets linked to higher or lower payoffs across two configurations. Participants relied exclusively on kinematic cues from hand shape changes in maximum grip aperture to infer their partner’s choices. There were two main findings. While participants preferred higher payoffs consistent with rational game-theoretic expectations, reliable kinematic cues overrode these expectations. When early grip aperture changes indicated the partner was reaching for a large target associated with a lower payoff, participants abandoned their default preference for higher payoffs. They chose the lower option instead, achieving a high coordination success rate. These findings demonstrate that people prioritize kinematic cues about others’ choices over theoretical assumptions about rational behavior when coordinating. This suggests that movement-based inferences about others’ actions in natural social interactions may be weighted more heavily than strategic reasoning when the two sources of information conflict.

## INTRODUCTION

Coordination is fundamental to human sociality. While many species cooperate and divide labor—from mammals and insects to plants and bacteria—humans are uniquely flexible in their social strategies. They can deploy multiple approaches to achieve the same goals, with simple actions offering choices: Selma might wait for Cameron to choose her glass or go first; Hayley might cook rice while Aysha fetches wine, or they might coordinate differently.

This flexibility, however, creates a puzzle. Despite decades of research, we still lack an adequate understanding of how humans coordinate so effectively. Game theory provides the most powerful analytical framework for studying such interactions (von Neumann & Morgenstern, 1944). A key insight is that coordination problems involve strategic decisions with multiple Nash equilibria—stable outcomes where each agent’s choice is a best response to others’ choices, and no one benefits from unilaterally changing strategy (Schelling, 1960; Kim, 1996). These best responses depend on expectations about others’ behavior, which may be based on direct observation (seeing you pick the large glass), but are often constructed from partial or conflicting information.

An important discovery of experimental game theory is that people coordinate smoothly without reliable information (Cooper & Weber, 2020). Sometimes they follow simple environmental cues: when sitting at a restaurant table, we typically drink from the glass in front of us, rather than from the glass in front of another guest. Physical proximity provides a salient focal point that requires minimal effort and cognitive processing. Similarly, if Hayley is a professional cook and Aysha is a sommelier, their obvious division of labor based on expertise likely generates a superior outcome.

Interactions of this kind have the structure of a so-called *HI-LO game*. The payoff structure reveals that one outcome (HI-HI) is superior because both parties prefer it to any alternative. While this is not the only equilibrium—choosing LO is *rationalizable* under specific belief configurations—, such beliefs seem intuitively odd. It would be rational for each agent to choose LO only if they expected the other to do likewise, but why would anyone hold such an expectation? Experiments confirm this intuition: most people choose HI when simultaneously playing this game without communication (Bacharach, 2006).

The cognitive mechanism behind this pattern remains unclear. Some game theorists have proposed a *Payoff Dominance Principle* (Harsanyi & Selten, 1988), which prescribes that agents choose the actions leading to the equilibrium everyone prefers. However, such a principle merely describes rather than explains the observed behavior. Other theorists suggest that the agents perceive Hi-Lo as a collective rather than an individual decision (Bacharach, 2006; Sugden, 2000; Colman & Gold, 2018). Since the outcome HI-HI is collectively superior to all alternatives, the agents should implement the action leading to the best collective outcome. A third proposal holds that the agents simulate each other’s behaviour, with each player seeking the best way to achieve the best outcome for themselves, rather than collectively (Morton, 2005; Guala, 2020).

Behind these theoretical contrasts lies substantial agreement about a core set of facts: when people lack information about others’ likely choices, they tend to choose HI, expect others to choose HI, and consider these expectations *rational* or justified.^1^ However, such information-poor conditions are relatively unusual in practice. Real-world interactions typically occur in informationally rich environments where agents receive various behavioral cues about others. In favorable circumstances, these cues are consistent and unequivocal. In other cases, they send conflicting signals that agents must resolve.

In the experiments reported here, we examine how human agents make rapid decisions like those required in a *HI-LO game* using partial, incomplete cues about their co-player’s moves. Our design builds on research showing that people predict others’ action outcomes through kinematic cues (Becchio et al., 2012, 2018; Krishnan-Barman et al., 2017; Scaliti et al., 2023). Ambrosini et al. (2011) found that observers reliably infer intended targets from hand preshaping movements, gazing at targets before the hand arrives (see also Ambrosini et al., 2012; Costantini et al., 2012a, 2012b). Ansuini et al. (2016) demonstrated that hand preshaping enables target prediction from the earliest phases of action observation. Our recent study combining kinematic analysis with machine learning revealed that people primarily use kinematic cues like maximum grip aperture (i.e., the maximum distance between index finger and thumb during reaching for grasping) for early-phase predictions, with target-specific predictive outcomes emerging over time (Fanghella et al., 2005).

In the present study, participants played with a supposedly remote partner, choosing their preferred payoff based on partial observation of their partner’s movements. In one configuration, the partner grasped a large ball for the higher payoff and a small one for the lower payoff; this mapping was reversed in another. Participants were unaware that the partner was virtual. We used validated video stimuli from our previous study (Fanghella et al., submitted)—motion capture recordings of grasping actions toward large and small balls, with movements depicted by red sticks connecting motion capture markers positioned on significant anatomical landmarks. Action clips were restricted to four time points (from movement start to, respectively, 10%, 20%, 30% or 40% of the reaching and grasping movement) to evaluate information processing requirements for coordination. Since no target object was visible, participants relied solely on hand shape changes over time to guess the partner’s moves.

We aim to determine whether participants modify their rational expectations about their partner’s moves based on kinematic information, specifically when kinematic cues provide sufficient evidence to change expectations that the partner will choose the higher payoff. If participants’ choices are determined primarily by the rational expectation that partners always choose the higher payoff, then early kinematic information should minimally influence their decisions. However, if kinematic information matters, participants’ choices should change once available cues identify the partner’s action outcome. When kinematic information indicates a lower payoff outcome, it should override the rational expectation that partners always choose higher payoffs.

## MATERIAL AND METHODS

### Action Execution

Kinematic recording occurred at the LAMoBiR (“Laboratorio di Analisi del Movimento e Bioingegneria della Riabilitazione”), IRCCS Fondazione Don Carlo Gnocchi. Two agents (a female and a male) were asked to reach for and grasp, with a whole prehension of their right (dominant) hand, two objects that differed in size (e.g., a large ball and a small ball, with diameters of 2 and 10 cm, respectively). At the beginning of each recording, the right hand was closed in a pinch and 70 cm from the middle of the distance between the two objects (i.e., 10 cm). The two objects (large and small) were randomly located on the left or right side, with the sides being counterbalanced for each target size. An acoustic signal informed the agent to start the movement. Retro-reflective markers were positioned on anatomical landmarks of their right hand and arm according to the same procedure described in Carpinella et al., 2006. The 3-D spatial coordinates of the retro-reflective markers were captured with an optoelectronic SMART system (B|T|S, Milan, Italy) and then analyzed using a custom Matlab program (The Mathworks, Natick, MA, USA), as in Piedimonte et al. (2015).

## High-Low Game Task

### Participants

Sixty-eight participants were enrolled in the experiment. Thirty-five participants were administered Configuration 1 (large ball associated with high payoff), and thirty-three participants were administered Configuration 2 (small ball associated with high payoff). Ten participants were excluded because of technical issues during the online testing sessions. The final sample comprised fifty-eight participants (Configuration 1: 29 participants, mean age 23.9, SD 6.8, 19 male; Configuration 2: 29 participants, mean age 24.4, SD 8.1, 14 male). A post-hoc sensitivity analysis revealed that our sample size was sufficient to detect a medium effect size (*pη*^2^ = 0.06) on the interaction of interest 2*2*4: Configuration (between-factor), Size and Time (within factors). We estimated a medium effect size for the interaction of interest based on a previous study (Fanghella et al., 2025), where the Size*Time within-factor interaction of interest (2*4) was large (*pη*^2^=0.81). The participants were naive to the purpose of the experiment, were right-handed, had normal or corrected-to-normal vision, and had no history of psychiatric or neurological disorders. The local Ethics Committee approved all research methods, which were carried out following the principles of the revised Helsinki Declaration (World Medical Association General Assembly, 2008). Written informed consent was obtained from all the participants

### Stimuli

The motion-capture recordings were edited to create video clips. Red sticks connecting the motion-capture markers presented a schematic upper limb and hand. The target objects were not given. The video clips were acquired at 40 Hz and lasted 2230 ms on average. The presentation plane was the functional plane where the grip movement occurs.

We randomly selected a subset of sixteen video clips, with a balanced representation of eight clips per action target. The selected clips differed in agent and target side (50% of videos for each agent and side). Frames from two example video clips are illustrated in Figure 1. The sixteen selected video clips were edited to represent four normalized time intervals (10%, 20%, 30%, 40%) of the movement time. The movement time was calculated by excluding the reaction time of the agent for each trial. Stimuli were validated in a previous study on early target prediction in human participants and machine learning algorithms (Fanghella et al., 2025).

**Figure 1.**
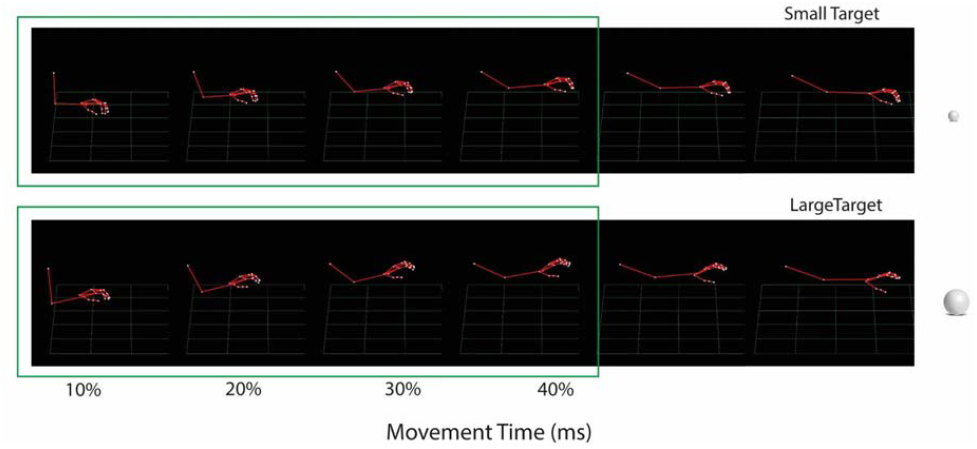
Example frames sampled from two acquired video streams at different times, one for the large ball and one for the small ball. Participants were shown 10%, 20%, 30%, or 40% of these videos. Note that the targets were not presented in the clips. (Two illustrative video clips are enclosed as supplementary movies.)

### Experimental design

All participants completed a High-Low game where they were asked to coordinate with a partner to obtain a monetary reward. Due to COVID-19 restrictions, the experiment was conducted online. The experimenter contacted each participant via Microsoft Teams and provided written and oral instructions. Specifically, the experimenter explained to the participants that they would play a High-Low coordination game online with a partner to gain a monetary reward. The participant was informed that the other player was connected in real-time from a kinematic lab to play the game together. However, unknown to the experimental subject, the other player was a virtual partner.

The experimenter explained to participants, who were assigned the role of Player 1, that their monetary payoffs depended on the combination of their choices and the choices of Player 2 (the virtual partner). Both Player 1 (the experimental subject) and Player 2 (the virtual partner) could choose either a high reward (6 euros) or a low reward (3 euros) for each round of the game. Player 1 was instructed that, if they chose the high reward and their partner made the same choice, they would both gain the corresponding payoff (6 euros), while if they both chose the low reward, they would both gain 3 euros; however, if they chose two different rewards (e.g., Player 1 chose the high reward, and Player 2 the low reward), both players would not gain any monetary payoff (0 euro).

Participants were asked to press as quickly as possible the ‘m’ or ‘n’ key (response keys were randomized across trials) to select either the high or low reward. During each game round, a matrix appeared on Player 1’s screen, displaying the payoffs associated with Player 1’s and Player 2’s choices (Figure 2). Specifically, Player 1 had to press the response key related to the reward of their choice and was told that Player 2 would have simultaneously selected the high or low reward by grabbing either a small ball or a big ball placed in front of their desk. The association between reward and target size was counterbalanced across participants in a between-group design (Configuration 1: large ball associated with high payoff; Configuration 2: small ball associated with high payoff).

**Figure 2.**
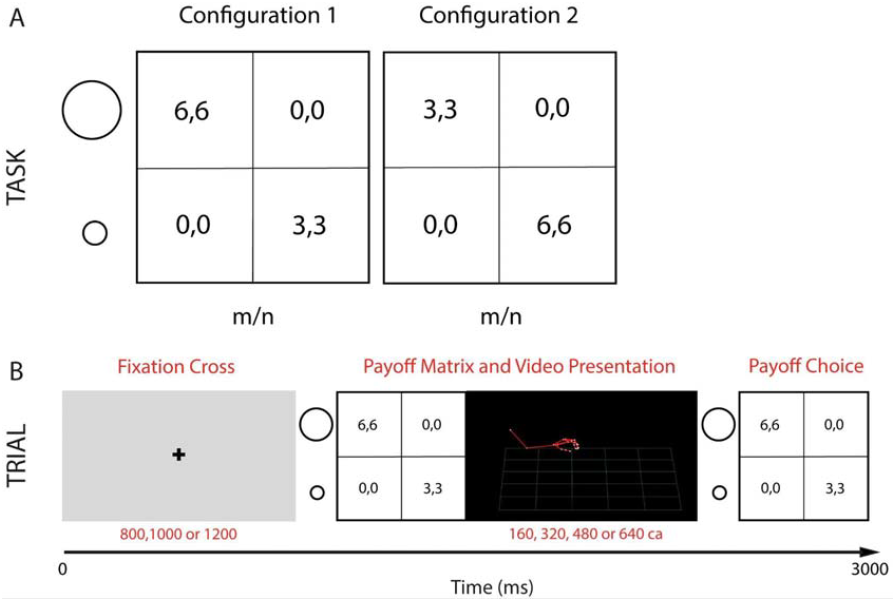
*Task*. A. In Configuration 1, for Player 2, a high reward (6) was always associated with the large ball, while in Configuration 2, Player 2’s high reward was always associated with the small ball. For Player 1, a high reward could be related to either button n or m in both Configurations. Participants could obtain a high reward (6,6) or a low reward (3,3) if they coordinated their choice based on the same payoff. However, if they selected two different payoffs, the monetary reward would be 0 for both players (0, 0). B. Structure of a trial. Each trial started with a fixation cross of variable duration. Together with the payoff matrix (A), participants observed a variable portion (10%, 20%, 30%, or 40% of the whole movement) of the kinematics of a grasping action toward a not-visible large or a small target, attributed to Player 2’s payoff choice. The response window, showing the payoff matrix, lasted in total 3000 ms.

Importantly, the experimenter explained to the participants that they would have the chance to guess their partner’s choice by observing a portion of a real-time reconstruction of the kinematics of Player 2 grasping action during each round of the game. This could help Player 1 guess Player 2’s payoff choice. The real-time reconstructions of Player 2 grasping actions, associated with payoff selection and displayed during each round, were actually pre-registered actions of two confederates, showing 10%, 20%, 30%, or 40% of a grasping action of either a small or a large ball, collected at an earlier stage (see the section *Stimuli*). The kinematics of Player 2’s movement was presented simultaneously to the payoff matrix to Player 1 (see Figure 2).

Before each experimental session, participants were asked to test whether the real-time reconstruction of Player 2’s grasping action of either the small or the large ball was effective. This trial session also aimed to test the compatibility between participants’ PC/laptops and the Eprime Go program used for the experiment by administering portions of the videos used during the testing session. Participants who dropped more than ten frames while videos were displayed were excluded from the final sample.

The experimental session consisted of 256 trials, grouped into four blocks (64 trials each). Before starting the experiment, participants completed a 16-trial practice. Each of the 16 target videos was presented 16 times, 4 for each movement time.

Every trial began with a fixation cross, lasting 800, 1000, or 1200 ms (variability was added to avoid habituation). Then, the configuration depicting the payoff matrix associated with the current trial appeared on the left, while the video clip depicting a portion of the grasping movement (10%, 20%, 30%, 40%), fictionally related to Player 2’s choice of payoff, was displayed on the right (Figure 2B). For Player 1, the response key associated with selecting the high reward was ‘m’ in 50% of trials and ‘n’ in the remaining 50% in randomized order. To counterbalance the association between large or small ball size and high or low reward, for virtual Player 2, grasping the large target was associated with choosing the high reward in Configuration 1, and grasping the small target was associated with choosing the high reward in Configuration 2. Each Configuration was administered to 50% of the participants (see section *Participants*). The configuration matrix was displayed for 3000 ms from its onset. Initially, it was presented next to the video clip depicting portions of the movement of Player 2, which had variable durations (approximately 160 (10% of action), 320 (20% of action), 480 (30% of action), or 640 ms (40% of action)). At the end of the video presentation, the configuration matrix remained displayed until 3000 ms from its onset. Participants’ responses were recorded during the whole 3000 ms time window. Before each new trial, participants were warned to get ready for a new round of the game.

All participants were emailed about their payoffs in the following days and received the corresponding monetary amount.

## Data Analysis

### Action Execution

A custom Matlab script was used to extract, for each considered time interval (10% to 40%), the max grip aperture, defined as the max distance between the markers of thumb and index during the reach-to-grasp movement (see Piedimonte et al., 2015), and max wrist velocity. As described in Fanghella et al. (submitted), a 2*4 repeated-measures ANOVA was run on grip aperture and wrist velocity for the 16 target videos with the within factors target Size (Large/Small) and Time (10%, 20%, 30%, 40%) of the whole movement.

### Hi-Low Game Task

#### Payoff Choice

First, we tested whether participants preferred a high payoff. We tested the overall *payoff choice*, calculating the percentage of high-payoff selections over the total amount of given responses, and we computed a one-sample t-test against zero to test if participants had a significant preference for high payoff.

To explore potential learning effects during the experiment, we compared the percentage of high-payoff choices during the first and last blocks with a repeated measures ANOVA with the within factor Block and the between-factor Configuration.. Finally, we analyzed how the rate of high payoff choice was modulated by size, time, and configuration through mixed repeated-measure ANOVAs. The factors of the ANOVA were: between-factor Configuration (1 or 2; 1: large ball associated with high payoff; 2: small ball associated with high payoff), and within-factors Size (Large/Small), and Time (10%, 20%, 30%, 40%).

#### Coordination

We focused on the trials in which Player 1 successfully coordinated with (virtual) Player 2 by choosing the same payoff. First, we measured the overall coordination rate and tested if the mean coordination was significantly above chance with a one-sample t-test against zero. Then, we analyzed how coordination strategies were modulated by size, time, and configuration through mixed repeated-measure ANOVAs in The factors of the ANOVA were: between-factor Configuration (1 or 2; 1: large ball associated with high payoff; 2: small ball associated with high payoff), and within-factors Size (Large/Small), and Time (10%, 20%, 30%, 40%).

Replicating previous analysis, we also investigated participants’ *Reaction Times* (RTs). Since participants could choose their payoff any time during stimulus presentation, RTs were calculated from the beginning of each video associated with the payoff matrix. Only RT for trials ending with successful coordination were considered in the analysis. The factors of the ANOVA were: Configuration (1 or 2), Size (Large/Small), and Time (10%, 20%, 30%, 40%).

In all statistical analyses, we applied Greenhouse-Geisser correction when appropriate to correct for violations of sphericity in F-tests (Keselman & Rogan, 1980). All post-hoc tests were carried out through Bonferroni-corrected pairwise comparisons. All statistical analyses were carried out in SPSS 29.0.1.0 and JASP 0.19.3.0.

## RESULTS

### Action Execution

#### Grip aperture

We found a main effect of Time (*F*_(3,18)_=121.147, *p* < 0.001, *pη*^2^ =0.953) and Size (*F*_(1,6)_=79.584, *p* < 0.001, *pη*;^2^ =0.930), as well as a statistically significant Size*Time interaction (*F*_(3,18)_=95.868, *p* < 0.001, *pη*;^2^ =0.941). Post-hoc test with Bonferroni correction revealed a greater grip aperture for the large object than for the small object from 20% up to 40% (mean±SE; 45.6±3.1 vs 30.3±1.1mm at 20% and 68.9±4.2 vs. 33.2±1.5mm at 30% and 84.8±4.6 vs. 35.3±1.8mm at 40%; *p* < 0.005 at 20% and *p* < 0.0005 at 30% and 40%). We did not find differences in grip aperture between the small and the large object at 10% of the time (28.8±1.7mm vs 26.8±0.5mm). These results are previously reported in Fanghella et al., submitted, and depicted in Figure 3.

**Figure 3.**
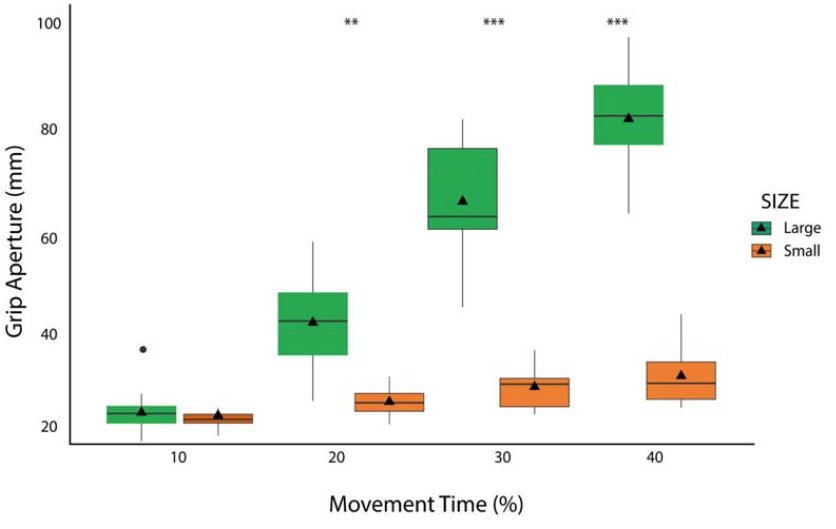
*Grip Aperture*. Boxplots with individual data points of the 16 selected kinematics videos for large and small objects. Results highlight significant differences in grip aperture at 20%, 30%, and 40% of the movement. \*\**p* < .01; \*\*\**p* < .00

#### Wrist Velocity

We found a main effect of Time (*F*_(1.652,9.913)_ =338.470, *p* < 0.0001, *pη*;^2^ =0.983). Conversely, no main effects or interactions involving the factor Size were significant (*p* > 0.05). Post-hoc comparison of the main effect of Time showed a significant difference in the velocity of the wrist between each time of movement (*p* < 0.005) except 30% compared to 40% time of movement (*p* =1) (unit mm/s; mean±SE; 10%: 271.4±10.4; 20%: 463.8±16.5; 30%: 569.3±18.0; 40%: 583.7±12.7).

### Hi-Low Game Task

#### Payoff Choice

The overall rate of high payoff choice was 58.61% of the total amount of given responses, regardless of coordination success. A one-sample t-test against zero confirmed that participants preferred the high payoff significantly above chance (t_57_ =25.37, p <.001). Results from the Configuration*Block repeated-measure ANOVA testing if learning effects occurred from first to last blocks revealed no significant main effect of Block (p =.199), revealing no overall changes in payoff choice strategies over time. However, we found a significant Configuration*Block interaction (F_(1, 56)_ = 14.147, p <.001, pη;2 =.202). Bonferroni-corrected post-hoc test revealed significantly reduced high-payoff choice in last block compared to first block when the high payoff was associated with the large ball (Configuration 1: first block, M = 61.07, SE = 3.228, last block M = 52.994, SE = 3,300, p =.004) but not when the high payoff was associated with the small ball (Configuration 2: first block, M = 57.438, SE = 3.228, last block M = 61.480, SE = 3.300, p =.524). The main effect of Configuration was not significant (p =.595).

We then analyzed the percentage of high payoff choices over all trials. The Configuration*Size*Time mixed-repeated measures ANOVA on high payoff choice yielded a Configuration*Size*Time significant interaction (F_(3, 168)_ = 29.971, p <.001, pη;2 =.349). Bonferroni-corrected post-hoc tests showed that in Configuration 1 there was a significant increase of high payoff choice when the ball depicted in the video was large compared to small at 10% (Large: M =50.152, SE = 4.715; Small: M = 44.465, SE = 5.343, p =.010), 20% (Large: M = 71.924, SE = 4.191; Small: M = 46.607, SE = 5.139, p <.001), 30% (Large: M = 78.660, SE = 4.783; Small: M = 48.219, SE = 5.181, p <.001) and 40 (Large: M = 79.544, SE = 4.842; Small: M = 46.544, SE = 5.312, p <.001). For Configuration 2, there was no significant difference in high payoff choice between small and large ball at 10% (Large: M = 74.503, SE = 4.715; Small: M = 76.586, SE = 5.343, p =.332); then, we observed a significantly enhanced high payoff choice for small compared to large ball at 20% (Large: M = 50.126, SE = 4.191; Small: M = 76.067, SE = 5.139, p <.001), 30% (Large: M = 33.447, SE = 4.789; Small: M = 68.056, SE = 5.181, p <.001) and 40% (Large: M = 28.649, SE = 4.842; Small: M = 66.130, SE = 5.312, p <.001).

The following interactions and main effects were also significant: Configuration*Size (F_(1, 56)_ = 41.679, p <.001, pη;2 =.427, Configuration*Time (F_(3, 168)_ = 38.928, p <.001, pη;2 =.410), and Time (F_(1.418 79.402)_ = 3.625, p =.046, pη;2 =.061). Post-hoc tests on the Configuration*Size interaction confirmed that, for each Configuration, participants chose the Size associated with the high payoff more often (Configuration 1: Large, M = 70.070, SE = 3.338, Small, M = 46.461, SE = 4.970, p <.001; Configuration 2: Large, M = 46.681, SE = 3.388; Small, M = 71.710, SE = 4.970, p <.001). Moreover, post-hoc tests on the Configuration*Time interaction revealed that participants chose less often the high reward in Configuration 1 compared to Configuration 2 at 10% (Configuration 1: Large, M = 47.308, SE = 4.925, Configuration 2, M= 75.544, SE = 4.925, p <.001), and they chose the high reward more often in Configuration 1 compared to Configuration 2 at 30% (Configuration 1: Large, M = 63.39, SE = 3.314, Configuration 2, M= 50.752, SE = 3.314, p =.009) and 40% (Configuration 1: Large, M = 63.049, SE = 3.349, Configuration 2, M = 47.389 SE = 3.349, p =.002). No differences were found at 20% (p = .458). Finally, post-hoc tests on the main effect of Time revealed that participants chose the high payoff more often at 20% compared to 40% of the time (20%: M = 61.181, SE = 2.562; 40%: 55.219, SE = 2.368, p =.009). All other post-hoc tests were not significant (all ps >.1).

#### Coordination

The overall coordination success rate, collapsing all factors, was 62.28%. A one-sample t-test against zero confirmed that successful coordination was significantly above chance (t_57_ =33.58, p <.001)

The mixed repeated-measures ANOVA on coordination scores yielded a significant Configuration*Size*Time interaction (F_(1.41, 81.54)_ = 3.790, p=.041, *pη*;2 =.063). Bonferroni corrected post-hoc pairwise comparison revealed that in Configuration 1, when the high reward was associated with the large ball, the level of coordination with the partner was higher when the video displayed Player 2 grasping a large ball compared to a small ball for the 20% (large ball: M = .719, SE = .042; small ball: M = .727, SE =.051; p=.014), 30% (large ball: M = .787, SE = .048; small ball: M = .527, SE =.051; p=.000) and 40 % (large ball: M = .794, SE = .048; small ball: M = .540, SE =.053; p=.000) movement times. Coordination for large versus small ball was not significantly different when movement time was 10% (large ball: M = .502, SE = .0.47; small ball: M = .556, SE =.053; p=.582).

We observed a different pattern in Configuration 2 (high reward associated with the small ball). Coordination was significantly higher when Player 2 chose the small ball compared to the large ball when the movement time was 10% (large ball: M = .255, SE = .0.47; small ball: M = .766, SE =.053; p=.000) and 20% (large ball: M = .499, SE = .0.42; small ball: M = .761, SE =.051; p=.001), while coordination did not differ between the large and small balls for the 30% (large ball: M = .666, SE = .0.48; small ball: M = .681, SE =.051; p=.818) and 40% (large ball: M = .714, SE = .0.48; small ball: M = .681, SE =.051; p=.440) movement times. These results are depicted in Figure 4.

**Figure 4.**
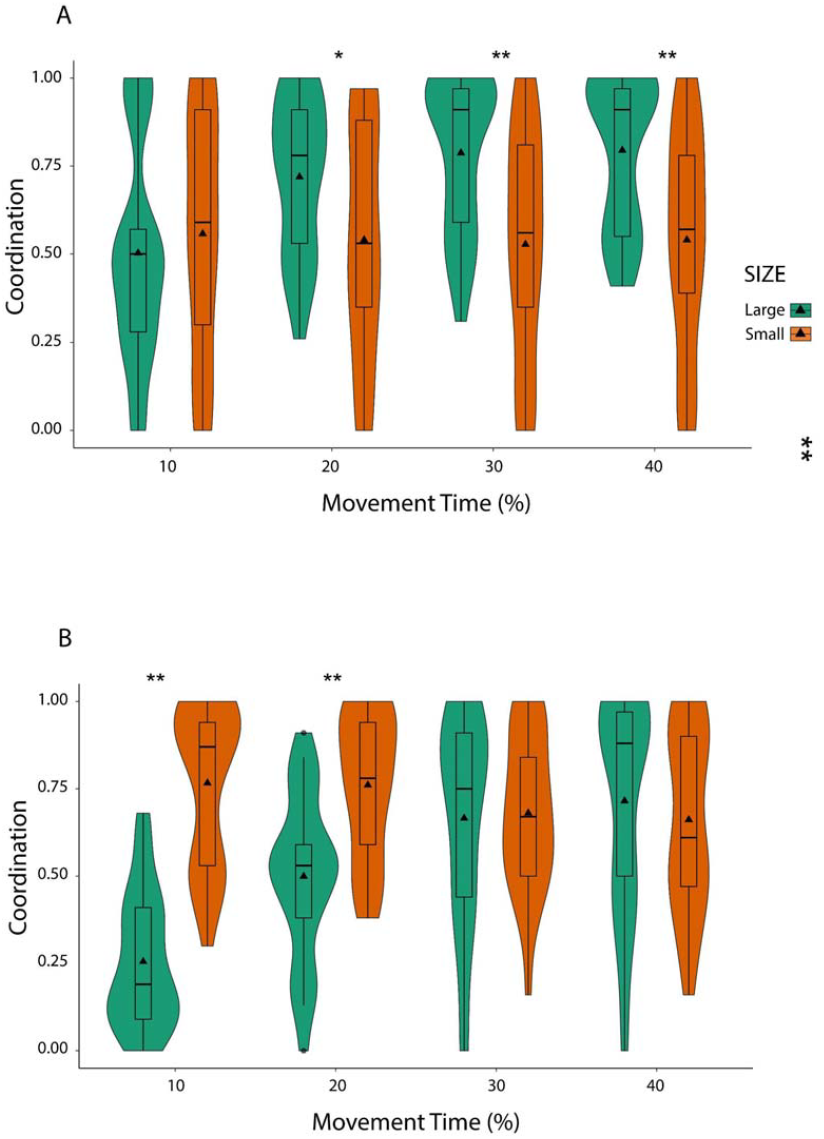
*Results: coordination*. A. Participants were more coordinated with the large compared to the small ball for 20%, 30%, and 40% times when the higher payoff was associated with the large ball (Configuration 1), and B. with the small compared to the large ball for the 10% and 20% times when the higher payoff was associated with the small ball (Configuration 2). *p<.05; **p<.01.

Moreover, we found a significant Configuration*Size interaction (F_(1, 56)_ = 13.641, p=.001, *pη*;2 =.196). As we expected, Bonferroni corrected post-hoc pairwise comparison showed higher coordination on the large ball compared to the small ball in Configuration 1 (large ball: M = .799, SE = .0.34; small ball: M = .540, SE = .049, p=.018), when the large ball was associated with the higher payoff, and higher coordination on the small ball compared to large in Configuration 2 (large ball: M = .533, SE = .0.34; small ball: M = .717, SE = .049, p=.007), when the small ball was associated with the higher payoff.

We also found a significant Size*Time interaction (F_(3, 56)_ = 38.365, p=.000, pη;2 =.407). Bonferroni corrected post-hoc pairwise comparison confirmed that overall participants were more coordinated when Player 2 grabbed the small ball after viewing the 10% time videos (Large ball: M = .378, SE = .0.33; small ball: M = .661, SE = .038; p=.000), while in the 30% (Large ball: M = .726, SE = .0.34; small ball: M = .604, SE = .036; p=.010) and 40% (large ball: M = .754, SE = .0.34; small ball: M = .600, SE = .037; p=.002) movement times, they were more coordinated when Player 2 chose the large ball. No differences were found in the 20% time (large ball: M = .609, SE = .030; small ball: M = .650, SE = .036; p=.427).

Finally, we found a significant main effect of Time (F_(1.69, 94.645)_ = 30.968, p=.000, *pη*;2 =.365), revealing lower coordination for the 10% time (M = .520, SE =.008) compared to the 20% time (M = .629, SE =.021, p=.000), the 30% time (M = .665, SE =.026, p=.000) and the 40% time (M = .677, SE =.027, p=.000). In the 40% time, participants were more coordinated than in the 20% time (p=.017).

The statistical analysis did not show a significant difference between coordination in Configuration 1 versus Configuration 2, as highlighted by the non-significant main effect of Configuration in the mixed repeated-measures ANOVA (Configuration 1: M = .620, SE = .026; Configuration 2: M = .625, SE = .026, p=.899). The main effect of Size was also not significant (large ball: M = .617, SE = .024; small ball: M = .629, SE = .035; p =.800).

#### Reaction times

The main ANOVA revealed a significant Size*Configuration interaction (F_(1, 50)_ = 5.935, p=.018, pη;2 =.106). This was explained by a significantly faster response for Large object in Configuration 1 compared to Configuration 2 (Configuration 1: M = 1149.517, SE = 74.180; Configuration 2: M = 1588.495, SE = 71.380; p = .042), but no differences across Configurations 1 and 2 for Small object (p = .410). Moreover, we found a significant main effect of Configuration (F_(1, 50)_ = 5.629, p=.022, pη;2 =.101), explained by overall faster responses during Configuration 1 compared to Configuration 2 (Configuration 1: M = 1395.598, SE = 73.976; Configuration 2: M = 1639.108, SE = 71.184). Moreover, we found a main effect of Size (F_(1, 50)_ = 60.070, p <.001, pη;2 =.546, revealing overall faster responses for the large ball (M = 1443.594, SE = 51.473) compared to the small ball (M = 1591.132, SE = 52.932), and a main effect of time (F_2.282, 114.106)_ = 8.541, p<.001, pη;2 =.146). The latter was explained by faster responses for the 10% movement time (M = 1449.060, SE = 48.165) compared to the 20% (M = 1511.679, SE = 47.616, p = .018), 30% (M = 1533.688, SE = 54.272, p = .041) and 40% (M = 1574.846, SE = 63.048, p = .001).

## DISCUSSION

This study investigated how humans coordinate using partial information about their co-player’s moves. Participants played a HI-LO game with a virtual partner, choosing their preferred payoff based on observing their partner’s grasping actions toward large and small targets. Partner’s hand movements were presented as video animations in which the hand is schematized by red sticks connecting motion capture markers positioned onto anatomical landmarks. In one configuration, the partner grasped a large ball for the higher payoff; in another, for the lower payoff. With invisible targets, participants relied solely on kinematic cues from hand shape changes to infer their partner’s moves.

There were two main findings. First, participants generally chose the higher payoff (HI) more frequently than the lower one (LO), regardless of their partner’s action outcome. Second, the coordination strategy differed across configurations. When the partner’s large ball grasping was associated with the HI choice, coordination was more successful when participants chose HI than LO at 20%, 30%, and 40% of movement time, but not at 10%. When the partner’s large ball grasping was associated with the LO choice, coordination was more successful when participants chose HI than LO at 10% and 20% of movement time, but showed no differences at 30% and 40%.

The first finding reflects a well-known pattern in game theory: in coordination games like HI-LO, players tend to choose HI, expecting others to do the same. Such an expectation may follow the *Payoff Dominance Principle* (Harsanyi & Selten, 1988), reflect collective decision-making where the chosen outcome benefits both players as a team (Bacharach, 2006; Sugden, 2000; Colman & Gold, 2018), or stem from simulating others’ possible choices (Morton, 2005; Guala, 2020). Whatever the underlying mechanism, players consider this expectation rational and justified.

The second finding reveals that kinematic information can override rational expectations when it provides evidence against HI choices, leading to successful coordination. In Configuration 2, participants abandoned their typical HI preference at 30% and 40% movement time, achieving high coordination success for both payoffs. In Configuration 1, participants also showed no HI preference at 10% movement time but achieved a very low coordination success rate (near chance level). This indicates that similar choice patterns produced different coordination outcomes depending on the experimental configuration and movement timing.

Prior research demonstrated that objects of different sizes could be identified through grasping kinematics. Observers are typically faster and more accurate in proactively gazing at objects when they can capitalize on hand preshaping suitable to the target (Ambrosini et al., 2011, 2012, 2015; Costantini et al., 2012a, 2012b). Similar effects occur in early infants: 6-, 8-, and 10-month-olds showed this pattern when observing an agent grasp or touch large or small objects (Ambrosini et al., 2013). Hand preshaping is critical for predicting action targets even in the earliest observation phases.

Ansuini et al. (2016) showed that target identification accuracy increased over time as kinematic information increased. Participants discriminated between large and small objects starting from 20% of the observed action, reaching maximum accuracy between 30% and 40%. Their performance mirrored relevant kinematic parameters, particularly maximum grip aperture (Ansuini et al. 2015). We recently found that prediction outcomes vary across target sizes over time. Participants were initially biased toward the small target (10% of movement time), perceiving minimal distance between index finger and thumb, with prediction rates remaining stable over time. In contrast, they predicted the large target from 20% of movement time onward, with prediction rates increasing as grip aperture increased. Machine learning algorithms corroborated this asymmetry, indicating that kinematic cues, rather than visual patterns, drive target size prediction (Fanghella et al., 2005).

This previous evidence provides a potential explanation for our second finding and the differences between Configuration 1 and 2. Participants successfully coordinated in Configuration 1 (where the large target is associated with the higher payoff) by choosing HI over LO starting from 20% of movement time, immediately achieving high success rates that remained stable at 30% and 40%. This result likely reflects the combined effect of participants’ rational expectation that partners would choose the higher payoff and their ability to identify the partner’s large target, given that kinematic information related to the grip aperture enabled target size discrimination from 20% onward. This could also explain why participants struggled to coordinate at 10% of movement time, when reliable kinematic information was absent—the perceptual bias interfered with rational expectations, reducing coordination success. The asymmetry between configurations at 10% movement time supports this explanation. In Configuration 2, where the small target yields the higher payoff, participants’ perception of small grip apertures aligns with their rational expectation that partners will choose HI. This consistency between perceptual evidence and strategic expectations produced a significantly higher coordination success rate at 10% movement time in Configuration 2, compared to Configuration 1, where the same perceptual cues conflicted with rational expectations.

In Configuration 2 (where the large target is associated with the lower payoff), participants also successfully coordinated by choosing the same payoff as their partner, without any substantial preference for HI over LO, at 30% and 40% of the movement time. That coordination success no longer depends on choosing HI over LO, as kinematic information about the partner’s grip aperture increases suggests that this information was sufficiently reliable to override participants’ rational expectation that players always choose the higher payoff. One might wonder why this doesn’t occur at 20% of movement time, given that kinematic analysis—particularly maximum grip aperture—enables discrimination of grasp types and target sizes at 20%, and people can identify potential targets with only 20% of action duration available.

A first explanation for this discrepancy is that, unlike previous studies, participants here had to modulate their payoff choice based on available information and target inferences, not simply infer potential targets. Given the increased task complexity, a more extended movement time (20% to 30%) may have been needed to infer target size with sufficient accuracy. A second explanation, partly compatible with the first, is that abandoning the deeply rooted expectation that rational agents always choose the higher payoff requires highly reliable contrary evidence. Kinematic analysis shows that maximum grip aperture increases significantly between 30% and 40% of movement time for large ball grasping (see Figure 3). Previous studies indicate that people’s ability to infer large targets increases notably around 30%, reaching ceiling levels maintained from 40% onward (Ansuini et al., 2016; Fanghella et al., submitted).

If this hypothesis is correct, Configuration 2 data confirm how deeply rooted the expectation is that rational agents always choose the higher payoff. They also indicate that not all correct kinematic inferences have equal reliability and that confidence in kinematic inference can be high even in early action phases. Ansuini et al. (2016) found that participants’ ability to recognize the correctness of their target size inferences increased with kinematic information. Our study corroborates this, further showing how increased kinematic information can boost confidence enough to change deeply rooted beliefs that rational agents always opt for the best solution.

That kinematic information can reveal essential aspects of decision-making in game theory contexts is not entirely novel. Turri et al. (2022) analyzed participants’ kinematics when responding in a motor version of the Ultimatum Game. They found that movements predicted their offers’ fairness and acceptance/rejection decisions. This indicates that choice-related movements reflect critical decision-making features, revealing ongoing social decision dynamics. Our findings demonstrate that people capitalize on early kinematic cues of others’ choice-related movements when deciding optimal choices for coordination, even revising deeply held beliefs, such as that rational agents always choose the best option.

Further research is needed to assess whether the interplay between rational expectations and kinematic information generalizes to different action types and strategic games. If so, this would provide more robust evidence of how people coordinate under partial and conflicting information conditions and prioritize different information types when making decisions. When this information concerns others’ actions, even very early kinematic cues can change deeply rooted beliefs about rational agent choices.

## Acknowledgments

This article was supported by the Department of Philosophy ‘Piero Martinetti’ of the University of Milan with the Project “Departments of Excellence 2018-2022” and “Departments of Excellence 2023-2027” awarded by the Italian Ministry of Education, University and Research (MIUR) (to MF, MTP, GB, FG, and CS), the PRIN 2022 grant “The extended hand: psychophysical and neural foundations of a robotic supernumerary finger’s use for grasping augmentation or recovery” (2022J72LFW_002 to CS), the PRIN 2022 grant “Motor resonance during action planning and social interactions: from single neurons to brain circuits” (2022SP5K99_002 to GB), and the PRIN 2022 grant (GPRIN202223FGUAL_01 to FG) “Normative Kinds: Values and Classificatory Decisions in Science and Policy-Making”.

## Author Contributions

Conceptualization, MF, CFC, FG., and CS; Methodology, MF, CFC, FD, GB, MR, and MFe; Formal analysis, MF, MR, and MFe; Investigation, MF, CFC, and MTP.; Writing –Original Draft, CS, MF, and FG; Writing – Review and Editing, CS, MF, CFC, FG, MR, and MFe.; Visualization, MF and CFC; Supervision, CS and FG; Funding Acquisition, CS and FG.

## Transparency and Openness

Data, Analysis code, and Research Materials can be found at https://osf.io/7wcpx/?view_only=591a7fc81a924c8e804614b25775a8eb.

## Footnotes

1 Whether it is *really* rational to have expectations of this kind is, however, controversial. Whenever we speak of rational expectations, in this paper, we refer to expectations that are perceived as rational or justified by the relevant agents.

